# EEG spectral exponent as a synthetic index for the longitudinal assessment of stroke recovery

**DOI:** 10.1101/2021.08.09.455664

**Authors:** J. Lanzone, M. Colombo, S. Sarasso, F. Zappasodi, M. Rosanova, M. Massimini, V. Di Lazzaro, G. Assenza

## Abstract

**Background:** Quantitative EEG (qEEG) can capture changes in brain activity that follow a stroke. Accordingly, EEG metrics could be used to monitor patients’ state and recovery. Although qEEG metrics traditionally focus on oscillatory activity, recent findings highlight the importance of aperiodic (power-law) structure in characterizing pathological brain states.

**Objective:** To assess neurophysiological impairment and recovery from mono-hemispheric stroke by means of the Spectral Exponent (SE), a metric that reflects EEG slowing and quantifies the power-law decay of the EEG Power Spectral Density (PSD). To relate neurophysiological recovery with patient’s functional outcome.

**Methods:** Eighteen patients (n=18) with Middle Cerebral Artery (MCA) ischaemic stroke were retrospectively enrolled for this study. Patients underwent EEG recording in the sub-acute phase (T0) and after 2 months of physical rehabilitation (T1). Sixteen healthy controls (HC; n=16) matched by age and sex were enrolled as a normative group. SE values and narrow-band PSD were estimated for each recording. We compared SE and band-power between patients and HC, and between the affected (AH) and unaffected hemisphere (UH) at T0 and T1 in patients.

**Results:** At T0, stroke patients showed significantly more negative SE values than HC (p=0.003), reflecting broad-band EEG slowing. Moreover, SE over the AH was consistently more negative compared to the UH and showed a renormalization at T1 in our patient sample. This SE renormalization significantly correlated with NIHSS improvement (R= 0.63, p=0.005).

**Conclusions:** SE is a reliable readout of the electric changes occurring in the brain after an ischaemic cortical lesion. Moreover, SE holds the promise to be a robust method to assess stroke impairment as well as to monitor and predict functional outcome.

## Introduction

Stroke is a leading cause of disability and constitutes a heavy burden for healthcare systems[1]. Monitoring brain activity with electroencephalography (EEG) could be a versatile instrument to aid the care of chronic and acute stroke patients. Indeed, given its ease of use, EEG is particularly suited for long-term monitoring of critical patients (i.e., intensive care units) as well as for the longitudinal assessment of patients with chronic stroke. In this context, quantitative EEG measures, such as ‘classical’ narrow-band Power Spectral Density (PSD), show a good correlation with stroke outcome [2–4] and the literature is rich of reports on how stroke affects the power of different EEG frequency bands [2, 3, 5]. Typically, the power of lower frequencies (delta, theta) increases, while the power of higher frequencies (alpha, beta) decreases[6]. However, narrow-band power measures (e.g. delta power) conflate the periodic and aperiodic features of the signal [7–9]. Indeed, band-power measures of the EEG do not take into account the aperiodic 1/f-like component[10]--that largely dominates the EEG PSD. This 1/f-like shape is a general property of brain dynamics, consistent across different spatial scales, from single neurons [11], to meso/macro scale oscillations[12–14], and is found in the decaying shape of the EEG’s PSD. The spectral exponent (SE) measures the steepness of the decay of the PSD background, relying on its 1/f-like structure. Specifically, it is estimated as the slope of the PSD, in log-log coordinates, once the bias due to oscillatory peaks is minimized[15–17].

The SE is closely related to the clinical concept of EEG slowing, given that a slower EEG entails a steeper PSD decay. Additionally, the ratio between slow and fast frequencies, often used as a quantitative measure for stroke assessment [2, 4, 5], is directly affected by underlying changes in the PSD decay. EEG slowing is commonly observed after stroke and has been known from a clinical perspective for a long time[18]. More recently EEG slowing has been described also in qEEG studies, showing increase in delta activity or delta/alpha ratio[3, 5]. EEG changes after stroke seem to be more prominent after large cortical ischemia[19] and have been related with the clinical status of the patient in different studies [20–23]. We propose that these EEG alterations could be also considered as the result of an overall change of the PSD shape, rather than changes in narrow-band power, thus justifying the use of a comprehensive measure such as the SE.

The slope of the PSD decay has also been linked to the neuronal balance between excitation and inhibition (E/I) [24]. As described by Gao and colleagues, when the contribution of the inhibitory population to local field potentials (LFP) was increased, the decay of the PSD in simulated local field potentials steepens [24]. This effect was confirmed in various experimental set-ups, studying the spatial and temporal modulations of the E/I in rodents, and under anesthesia in macaques. Furthermore, the PSD decay in humans resulted to be steeper in conditions typically related with increased inhibition, such as NREM sleep [10, 25, 26] or general anesthesia [17], when compared to wakefulness – as indexed by more negative SE values [25, 27–29]. Therefore, the use of SE is gaining popularity also in the study of neurologic conditions [30–33]

Interestingly, more negative SE values (paralleled by reduced E/I ratio) were found over the affected hemisphere in rat models of stroke[34]. Since the SE captures EEG slowing, and is sensitive to alterations in the E/I balance, we expect it to be a reliable measure of the neurophysiological alterations in stroke. Accordingly, shifts in E/I ratio are a paramount feature in the neurophysiology of stroke[35] and several studies using Transcranial Magnetic Stimulation (TMS) consistently show altered excitability over the affected hemisphere [36, 37].

Given the above, we hypothesize that the SE, an index of the broad-band aperiodic EEG activity, could be a good estimate of the state of cortical circuits in stroke patients, and be predictive of post-stroke functional outcome.

Here we test the SE on a longitudinal EEG data set and assess its sensitivity to the effects of both acute and chronic stroke, and its modulation following one month of standard physical rehabilitation. We show that the SE of EEG is a good read-out of ischaemic brain lesions as it is intrinsically linked to the neurophysiological fingerprint of stroke.

## Materials and Methods

For the purpose of this study, we retrospectively analysed EEG recordings from 23 patients (14 Males, age 72±9.5 y.o. mean±sd, 22/23 right-handed) diagnosed with mono-hemispheric stroke (14/23 left hemisphere stroke, 13/23 strokes with cortical involvement) in the territory of the middle cerebral artery (MCA) and 16 Healthy Controls (HC) matched for age and sex (7/16 Males, age 68±10 y.o. mean±sd, 15/16 right-handed). EEG were recorded at Campus Bio-medico University of Rome and “Casa di cura San Raffaele Cassino”. Each patient had EEG recorded at T0 in the acute phase (a median of 6 days, Inter Quartile Range (IQR) 4|10 days, after the event), and at T1 after two months, at the end of rehabilitation (median 77 days, IQR 62|88 days after the event).

Between T0 and T1 time points all patients underwent a standardized, 1 month long, protocol of rehabilitation based on physical therapy. EEG recordings consisted of 10 min resting, waking EEG with eyes closed.

Clinical inclusion criteria were: **i)** First ever ischemic stroke of MCA territory confirmed by MRI; ii) Evidence of motor/sensory deficit of the upper limb as assessed by a Neurologist.

Exclusion criteria were: **i)** clinical history of previous stroke. **ii)** If patients could not comply with EEG recording. **iii)** If there was clinical/radiological evidence of acute bilateral involvement, brain haemorrhage, dementia or other neurodegenerative diseases such as Parkinson’s disease.

Exclusion criteria from EEG analysis were: 1) more than 1 bad channel (out of 19) in the EEG recording 2) lack of at least 180 seconds free from artefacts (jumps in the EEG signal, head movement, electrode pop) 3) lack of at least 180 seconds of closed eyes wakefulness.

If any of said criteria occurred in the EEG at T0 or at T1 the patient was excluded from the study.

At both T0 and T1 neurological status was assessed using the National Institute of Health Stroke Scale (NIHSS)[34]. NIHSS is an 11 items scale used in the assessment of clinical impairment related with stroke, range 0-42, with higher values reflecting more severe damages. Patients underwent 1.5 Tesla MRI scan at T0 as part of the diagnostic work-up, and the affected hemisphere was determined according to cerebral imaging and clinical findings. Lesion locations were subdivided into cortical (cortical or cortical/subcortical involvement) and purely sub-cortical (no clinical/radiologic evidence of cortical involvement) according to MRI findings. Supplementary Table 1 shows the NIHSS scores and the location of the ischaemic lesion in our patients.

### EEG recordings

Both groups of MCA patients and HC underwent 19 channels EEG recording with standard 10-20 montage[38]. 10 minutes of eyes closed EEG were recorded. Since the SE is minimally influenced by local spectral peaks (e.g. alpha peak) [17], the eyes closed condition was chosen given that it helps minimizing eye movements, as well as scalp muscle artifacts during the recording. The awake state of subjects was continuously monitored by EEG inspection. Signals were recorded with a 32 channel Micromed system (SystemPlus software; Micromed, Mogliano Veneto, IT) sampled at 256 Hz (16 bit A/D conversion) and online referenced to an electrode placed on the digitally linked mastoid. Contact impedance was kept below 5KΩ. Data was exported in EDF format for further analysis.

### Pipeline to analyze the EEG signal

EEG recordings were analyzed using MATLAB^©^ native code and the FieldTrip toolbox [39]. The pipeline for the analysis focused on eliminating the major confounders of SE. On the one hand, eye movements and EEG signal jumps increase delta power causing an artificially steeper slope, and on the other hand, intense muscular activity creates a bump in the high frequency range, causing an artificially shallower slope.

EEG were imported from EDF format with acquisition reference, channel location was added to the EEG structure. Long-range linear trends in the time-series were removed. Outliers among EEG channels were recognized using Z-score deviation over channels; the rejection threshold was set by visual inspection of each recording. Single bad channels were interpolated (nearest neighbour). Data was filtered with an IIR high-pass (5^th^ order Butterworth filter with a 0.5 Hz cut-off) and a notch filter centered at 50 Hz. Scalp muscle artifacts were removed using Canonical Correlation Analysis (CCA), as described by De Clercq et al [40]. Temporal autocorrelations in the EEG components were calculated with a sliding window of 10 second and 1 data point delay (1/256 Hz = 3.9 ms), components were ordered from the most autocorrelated to the least autocorrelated. Components with low autocorrelation (and thus likely of muscular origin) were removed from each window. The rejection threshold for autocorrelation was set at 0.7; this threshold was heuristically determined by inspection of time-series, PSD and scalp topography of each component. Data were then re-referenced to average reference. Independent Component Analysis (ICA) was performed and only components with clear ocular artifact were rejected by visual inspection of component’s topography, time-frequency and time series (average number of rejected components across EEG recordings: 1±0.7). Signals were visually reviewed and residual eye movements or EEG jumps were excluded from PSD calculation. The PSD was estimated using Welch’s method (2 seconds window, 50% overlap). SE of the 1-40 Hz-range was estimated for each pre-processed EEG channel. The code to estimate the SE is openly available online (https://github.com/milecombo/spectralExponent) and thoroughly explained in Colombo et al [26]. The topography of the SE value for each channel was plotted for visualization. In Fig.1 we show a representative process of SE calculation, further insight in SE depending on the frequency window is shown in Supp. Fig.1. Delta (1-4 Hz), Theta (4-8 Hz), Alpha (8-12 Hz) and Beta (13-20 Hz) narrow-band power was also calculated for each EEG channel in order to compare SE with other quantitative measures derived from the PSD. Specifically, for each frequency band we computed the absolute band power as well as the normalized relative power (band power/total power). In addition, we computed Delta/Alpha ratio and Theta/Alpha ratio given their common application in the EEG stroke evaluation[41].

**Fig.1.**
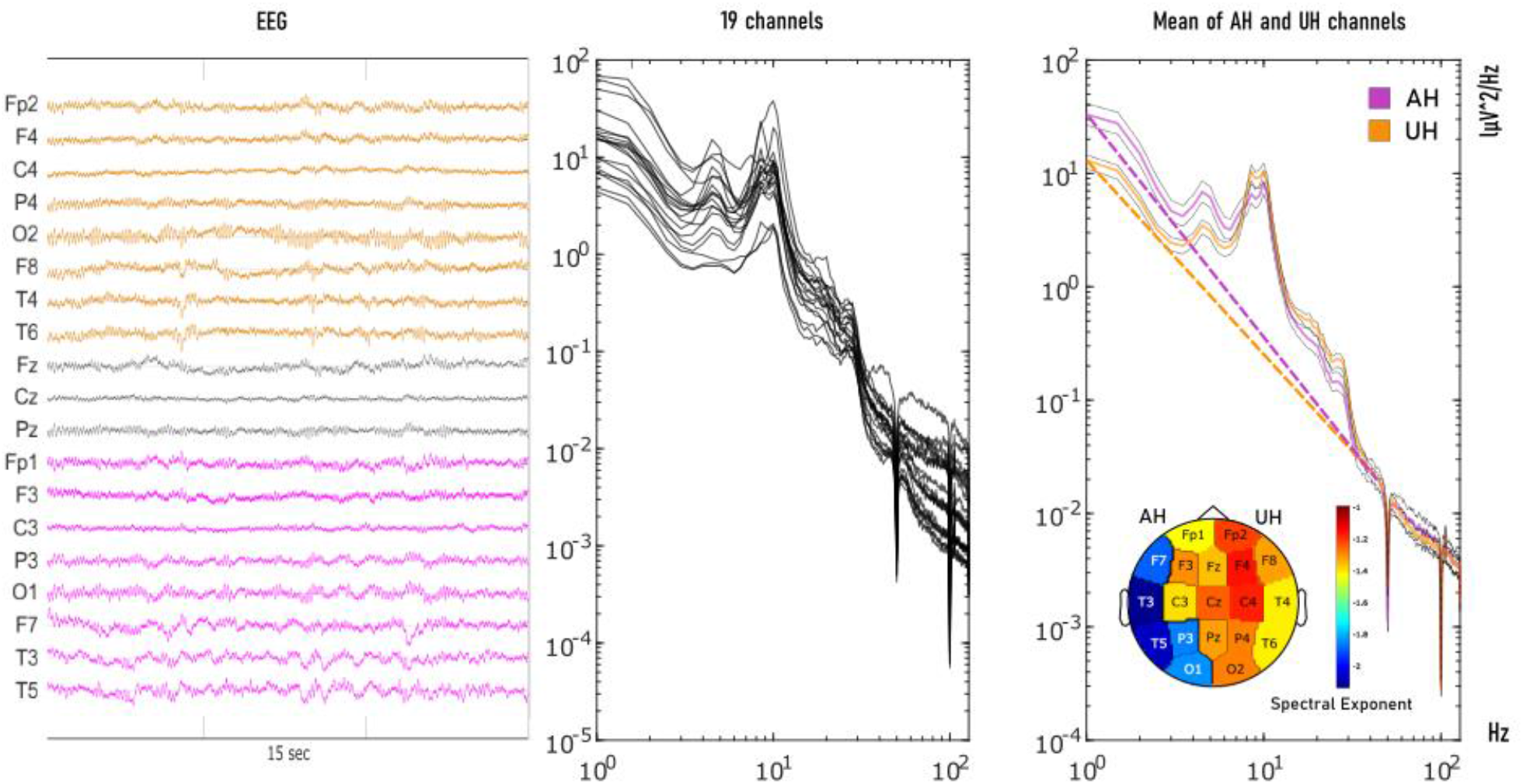
The spectral exponent reflects the decay of the PSD across a broad range of frequencies. Left panel: representative 15-s EEG traces for one patient (patient #17; left parietal stroke) recorded at T0. All channels with average reference are shown (AH in purple; UH in orange). Middle panel: PSD of the same EEG recording estimated (Welch) for all 19 channels after average referencing. Right panel: average PSD across AH channels (Purple), UH channels (Orange). The dashed lines represent the power-law fitted to the 1-40 Hz frequency range and the slope of these lines corresponds to the SE value. In the lower right corner we show topographic representation of the SE values in each channel.

### Statistical analysis

The SE of PSD in the 1-40 Hz range results in a single value for each EEG channel. Data normality was checked using a quantile-quantile method. To test overall difference in SE between stroke patients and HC, we first compared the between-groups average SE across the 19 channels in each EEG recording at T0 and at T1, using unpaired t-test.

Then, we assessed the inter-hemispheric effect of the lesion on SE for the stroke patient group. We discarded midline EEG channels (‘Fz’;’Cz’;’Pz’) and sorted the remaining 16 channels into the affected and unaffected hemisphere (AH and UH) for each patient individually, according to neuroimaging evaluation and clinical features. The mean SE value of the AH and of the UH was calculated. A similar approach was used for HC to assess SE differences between the right and the left hemisphere.

We hereby employ four condition labels, to index both the lesion-side and the time point: AHT0 (affected hemisphere, time point 0); UHT0 (unaffected hemisphere, time point 0); AHT1 (affected hemisphere, time point 1); UHT1 (unaffected hemisphere, time point 1). A general linear model was built in R^©^[42] in order to assess the main effects and their interactions. Specifically, a two-way repeated measures ANOVA was built, with the SE as dependent variable, hemisphere as within-subjects factor (levels: AH and UH), and time as within-subject repetition factor (levels: T0 and T1). Greenhouse-Geisser correction was used in the event of sphericity violation. Post-Hoc tests were performed using FDR correction for multiple comparisons. ANOVA assumptions were verified by quantile-quantile plotting and residual plot.

In addition, to verify the recovery of inter-hemispheric differences, the difference between the SE of the AH and UH was calculated for both T0 and T1 (SE[AH]- SE[UH]) for each patient, and a t-test against 0-mean assessed its change from T0 to T1.

The role of cortical vs. subcortical lesions on SE was further assessed by means of unpaired t-test, comparing patients with cortical vs purely subcortical lesions.

Pearson’s correlation of the NIHSS scores was calculated with the SE values in the AH and UH, as well as with the AH-UH difference, for T0 and T1. In addition, we computed the effective recovery rate (ER)[43], a dynamic measure of patients’ functional recovery, and also assessed its correlation with the SE at both timepoints. ER is calculated as the percentage of the occurred improvement with respect to the total possible improvement, taking into account that NIHSS = 0 corresponds to the absence of clinical symptoms:

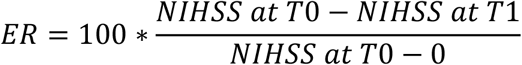

Finally, we performed an exploratory analysis of other narrow-band PSD metrics: we tested differences between AH and UH and T0 and T1 in absolute power and normalized relative power for all frequency bands, as well as in Delta/Alpha and Theta/Alpha ratios using Wilcoxon signed-rank tests. These exploratory comparisons are presented without any form of correction in order to maximize sensitivity. SE exponent statistics are reported as mean±standard deviation (sd). PSD metrics data are reported as median with Inter Quartile Range (IQR). For all statistical comparisons, the alpha level of significance was set at 0.05.

## Results

### Study population

5 patients matched the exclusion criteria due to poor compliance to EEG recording and/or poor quality of EEG signal and were thus not included. After removing these patients, our population comprised 18 patients (mean age 71.8±8.9 years, 10 males, 17 right-handed, 11 left hemisphere stroke, 12 with cortical involvement) and 16 Healthy Controls (HC) participants (7/16 Males, age 68±10 y.o. mean±sd, 15/16 right-handed).

### Clinical features

None of the patient enrolled underwent systemic thrombolysis, since they arrived at the emergency room out of time, according to the latest guidelines for stroke management[44]. NIHSS scores at T0 (median 5, IQR 2|9) were significantly higher (T(19)=2,54 p=0.02) than at T1 (median 2, IQR 0.25|2). This functional recovery was also quantified by means of the ER (median 0.634, IQR 0.34|0.96).

### Spectral Exponent in stroke patients vs healthy controls

Patients at T0 had a significantly steeper PSD (more negative SE values) compared to HC subjects (comparisons were made on the mean of all 19 channels from each subject),

**HC:** SE:−1.12± 0.24, **PatientsT0:** SE:−1.48±0.27 mean± sd; **T test**: T= −3.24, p=0.0029).

The same comparison for patients at T1 was only marginally significant (**PatientsT1:** SE −1.33±0.30; mean± sd; **T test** :T= −1.54 p=0.05), suggesting an overall renormalization of SE values over time in our patients group. As expected, HC did not show any significant inter-hemispheric asymmetry in the SE (**SE-Left** −1.13±0.22 mean± sd; **SE-Right**=−1.10±0.25 mean± sd; **T test:**T = −0.28, p-value = 0.77).

### Effects of lesions and time on SE in stroke patients

Figure 2 displays the two-way repeated measures ANOVA which showed significant main effects of both time (F [1,17] = 4.77, p=0.043) and hemisphere (F [1,17] =16.88 p=0.001) in the patients’ group. A significant interaction between time and hemisphere factors was also found (F [1,17] =18.64, p=0.0004).

**Fig.2.**
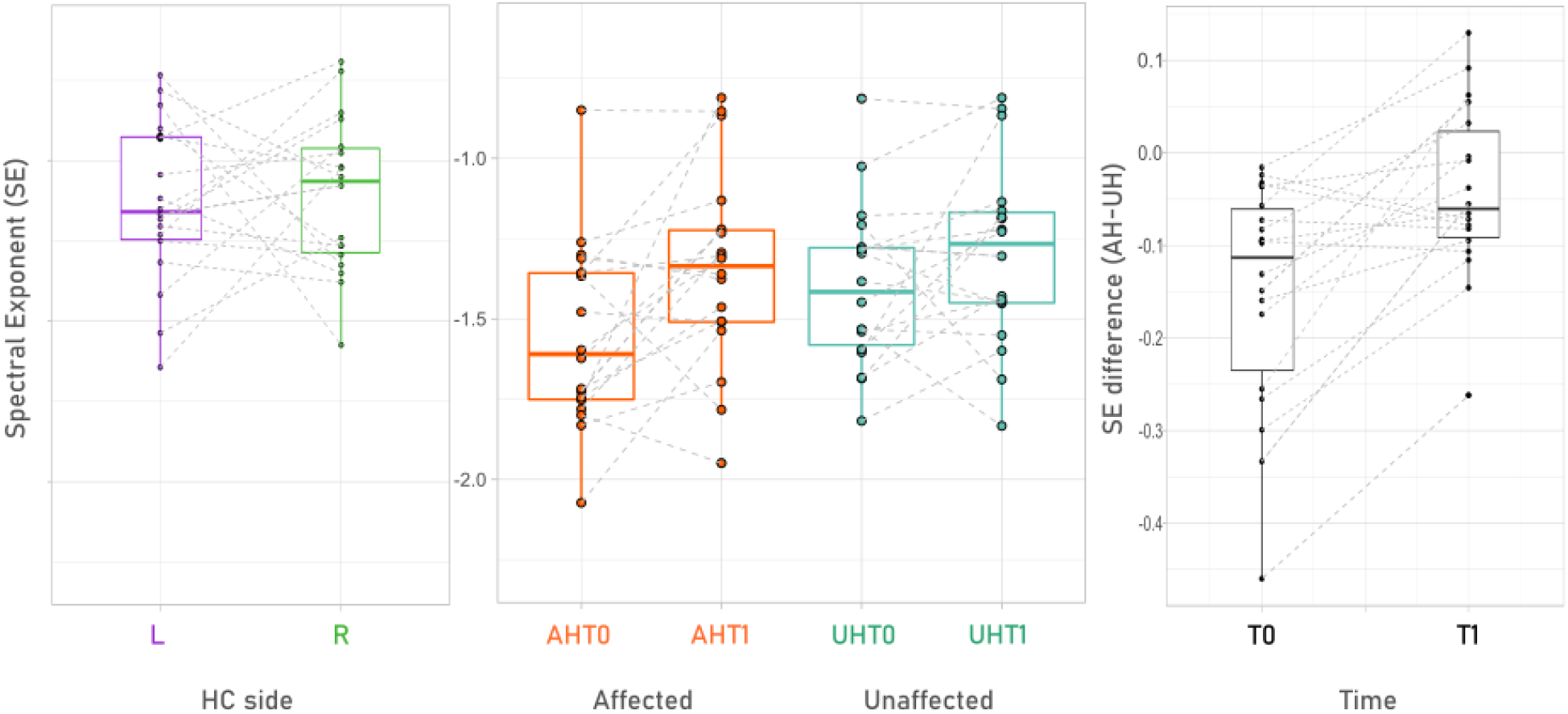
The SE is more negative on the affected as compared to the unaffected hemisphere; the SE interhemispheric difference decreases over time, renormalizing to the level of healthy controls. In the middle we show SE values for the affected (AH, orange) and unaffected (UH, blue) hemisphere at T0 and T1, as boxplots with overlaying case distribution. Dotted lines connect T0 and T1 of each patient. Healthy controls (HC) SE values from left (L, purple) and right (R, green) hemisphere are shown on the left side, with the same scale. On the right side we show how the difference between SE in the AH and UH changes over time.

Post-hoc analysis (FDR-corrected for multiple comparisons) evidenced significant SE differences in the following comparisons.

Inter-hemispheric differences at T0:

AHT0-SE:−1.55±0.289 vs UHT0 SE-1.40 ± 0.254, mean± sd; T= 5.19 p=0.0001);

Change in the affected hemisphere over time:

AHT0-SE:−1.55±0.289 vs AHT1 SE:−1.34 ± 0.309, mean± sd; T=2.74 p=0.003);

Affected hemisphere at T0 against its contralateral at T1:

AHT0-SE:−1.55±0.289 vs UHT1 SE:−1.30 ± 0.285, mean± sd; T=3.41 p=0.001).

Overall, these results show the occurrence of a steeper PSD slope over the affected hemisphere that renormalizes over time. These findings are further supported by the significant reduction in the inter-hemispheric difference (SE[AH]- SE[UH]) between T0 and T1. (**T0:** −0.1522±0.1244, **T1**:−0.0567±0.1243 mean± sd; **T test**: T=−3.6847, p=0.0018).

The type of lesion did not affect changes in the SE, since we did not find significant differences in SE when comparing SE between cortical (12 subjects) and non-cortical (6 subjects) lesions at T0 and at T1, both in the AH and in the UH (all P >.175 after correction).

### SE correlation with clinical indicators

Exploratory correlations of the SE with the NIHSS clinical scale and ER were performed across time-points in the affected and unaffected hemispheres. Results are shown in Table.1.

**Table.1.** Spectral exponent allows to predict functional recovery. The table shows correlation between SE values and clinical scores (un-corrected p-values). Significant values are highlighted in grey, significant findings after Bonferroni correction are highlighted in light blue.

|  | NIHSS T0 | NIHSS T1 | ER |
| --- | --- | --- | --- |
| **AHT0** | $r=-0,307$; $p=n.s.$ | $r=-0,478$; $p=0,04$ | $r=0,630$; $p=0,004$ |
| **UHT0** | $r=-0,255$; $p=n.s.$ | $r=-0,412$; $p=n.s.$ | $r=0,602$; $p=0,008$ |
| **AHT1** | $r=-0,482$; $p=0,04$ | $r=-0,707$; $p=0,001$ | $r=0,582$; $p=0,01$ |
| **UHT1** | $r=-0,436$; $p=n.s.$ | $r=-0,591$; $p=0,01$ | $r=0,509$; $p=0,03$ |

The SE showed good correlation with clinical outcomes. SE values at T0 showed less degree of correlation with NIHSS (both T0 and T1) compared to values of SE at T1. Meanwhile, ER was always correlated with SE regardless of the hemisphere or timepoint, with the strongest correlation being with SE in the AH at T0. In Fig.3 and Fig.4 we show the relation between SE and clinical scores, evidencing a similar pattern across cortical and non-cortical lesions.

**Fig.3.**
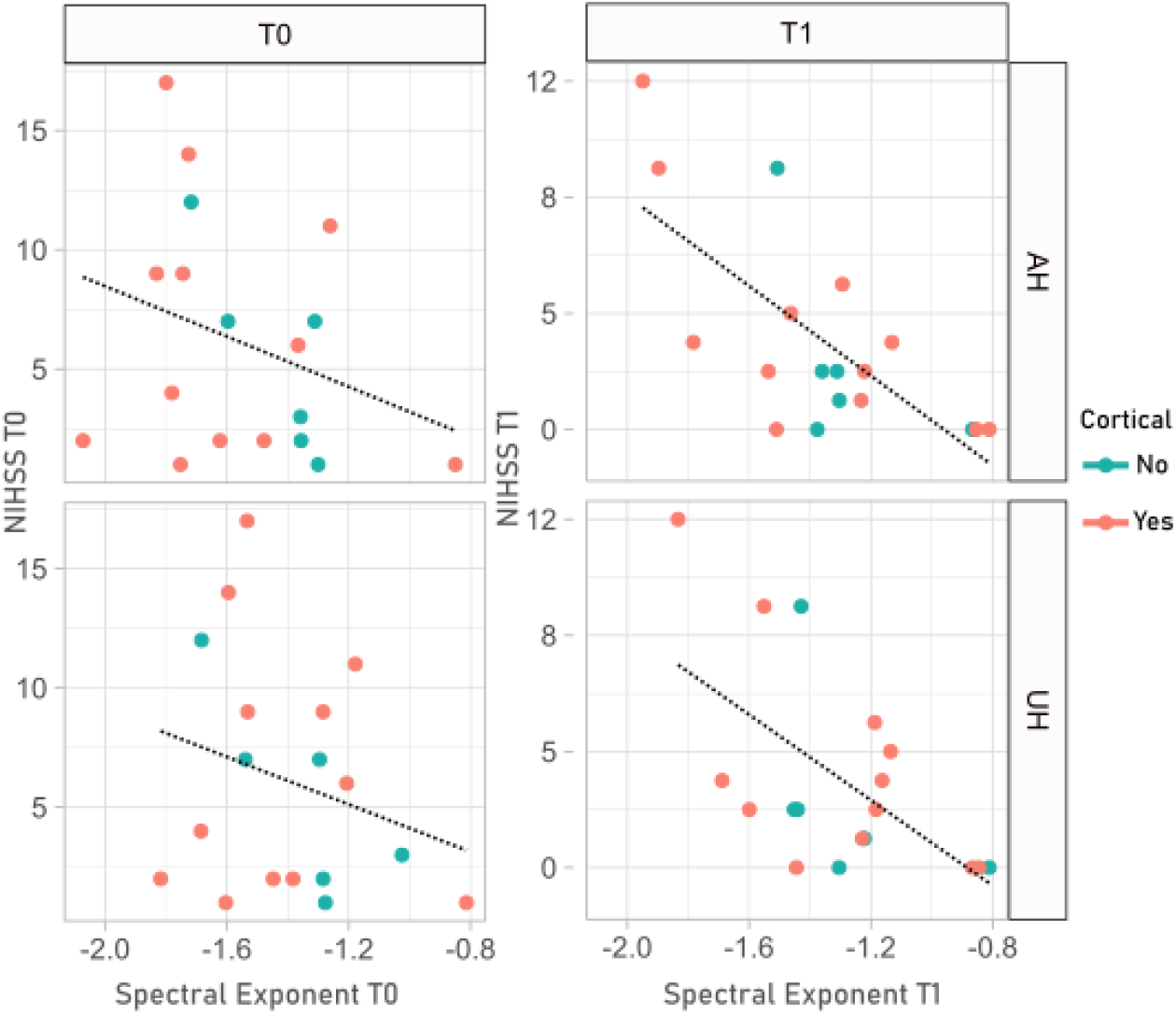
The SE is correlated with clinical current status. The figure shows the correlation between SE values in the AH and UH and NIHSS, here we show the relation of SE with concomitant values of NIHSS (T0 with T0, T1 with T1). Lesions with subcortical or cortical involvement are colour coded.

**Fig.4.**
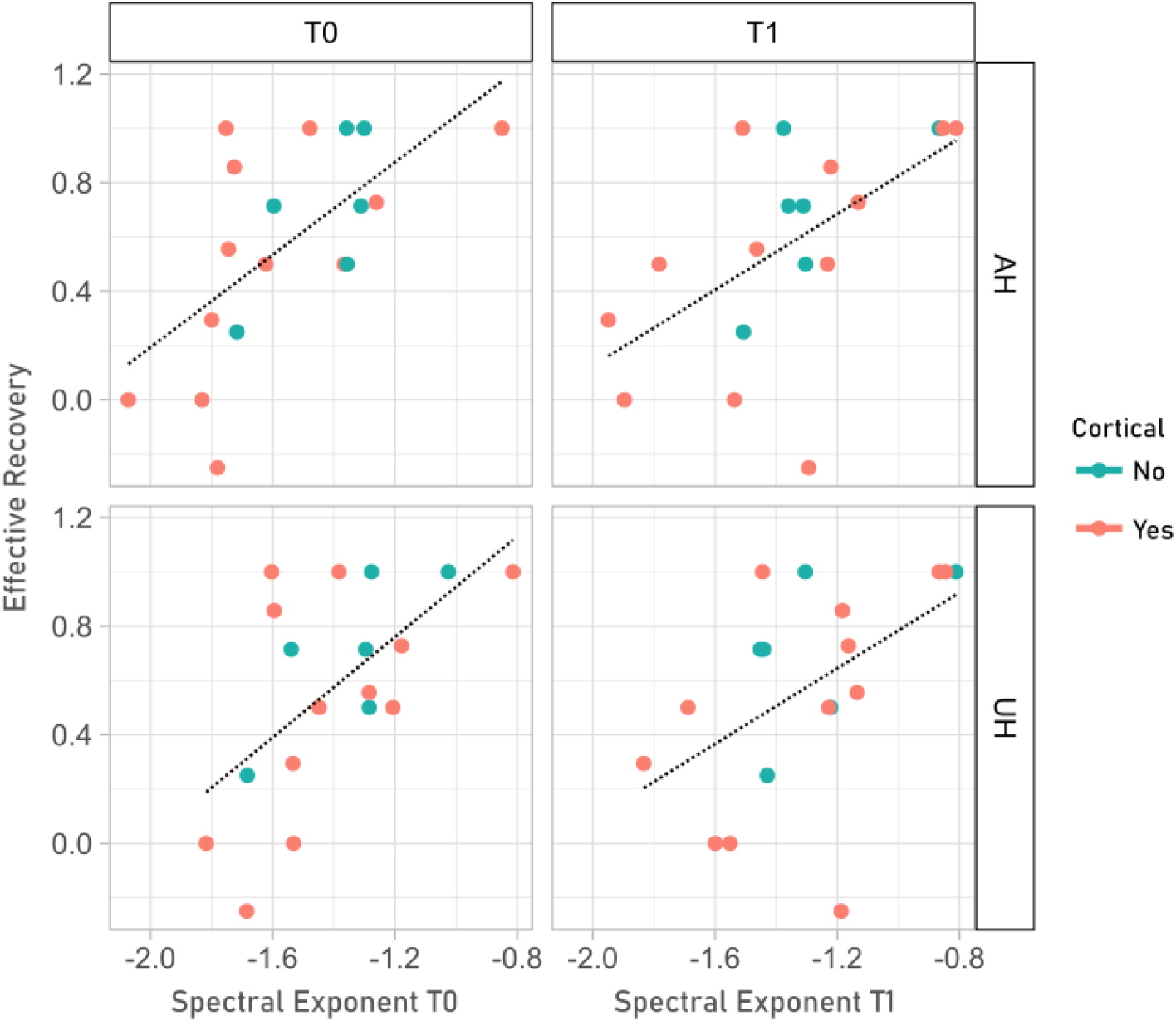
The SE allows predicting post-stroke functional recovery. The figure shows the correlation between SE values, in different hemispheres and across time points, with the effective recovery (ER) of patients. Lesions with and without cortical involvement are color coded.

### Conventional narrow-band power metrics

In Supplementary Table.2 we show the results of multiple tests comparing differences between conventional narrowband metrics derived from the PSD: absolute power, relative power as well as Delta/Alpha ratio and Theta/Alpha ratio. Commonly used spectral indexes did not evidence differences between the AH and the UH, showing only trends for Delta power and Delta/Alpha ratio (P>0.14). Significant differences (P<0.05) between T0 and T1 PSD were evidenced, especially when normalized measures were used, such as relative power or Delta/Alpha ratio. Specifically, higher Relative Delta power was evident at T0 both in the AH and UH, while Relative Alpha and Relative Beta bands were found lower at T0 both over the AH and the UH. Also, Delta/Alpha ratio was significantly higher at T0 compared to T1.

## DISCUSSION

In this study, the spectral exponent identified the lesioned hemisphere in patients affected by MCA stroke, and its longitudinal changes paralleled the recovery of clinical impairment. Specifically, we showed that mono-hemispheric stroke is characterized by a steeper PSD decay, indexed by more negative global SE values, compared to HC, shortly after the insult (T0). Most importantly, the SE was associated with the side of injury, with invariably more negative values over the affected hemisphere compared to the contralesional one, across all patients. Further, we showed that due to a prominent SE renormalization over the affected hemisphere, both local and global SE values attained similar values to those of HC after two months (T1). Highlighting the clinical relevance of these findings, the degree of SE renormalization reflected the degree of functional recovery. Overall, this study proposes the SE as a reliable neurophysiological fingerprint of stroke.

### The SE as a broad-band synthetic index

Previous literature on qEEG in stroke proved that after brain ischemia a significant change in Delta band spectral activity is recorded [41, 45, 46], particularly in cases with large cortical involvement [19]. Additionally, many studies described a close concordance between the area of maximum Delta power and the ischaemic area as shown by brain imaging (CT and MRI), suggesting that Delta activity might localize the injured area[41, 47–51].

Further improvement on the specificity of qEEG was warranted by the introduction of the Delta/Alpha ratio (DAR), which in many cases showed better clinical correlation than Delta power alone [2–5]. The DAR, measuring the ratio between slow and fast oscillations, grossly reflects the overall PSD shape, which is quantified by the spectral exponent. However, the DAR might be vulnerable to increases or decreases of Alpha/Delta spectral peaks such as alpha activity in the occipital derivation (eyes open vs closed) or increased delta activity due to non-cerebral artifacts (eye movements). Moreover, in our population the DAR was surpassed by the SE.

We here propose the SE as a novel qEEG synthetic metric, able to evidence clinically relevant asymmetries in brain activity, while providing at the same time a robust and comprehensive representation of the PSD.

### Neurophysiological determinants of the SE: E/I balance

Other than predicting functional outcome, the SE might help to better understand the pathophysiologic consequences of focal brain injury, by indexing alterations of the E/I balance.

Indeed, after ischaemic stroke the E/I balance in the lesioned area undergoes significant changes [52, 53] that indirectly reflect on the EEG [54]. During the hyperacute phase (first hours) neuronal depolarization and glutamate release cause transient hyperexcitability, thus increasing the E/I ratio [55, 56]. Thereafter, during the sub-acute and chronic phases (after cell death has transpired), GABA currents are enhanced [57], causing a decrease in excitability with lower E/I ratio [58]. These changes are thought to reflect on the EEG with increased low frequency activity and decreased fast frequency activity, which results in a steeper PSD shape (more negative SE values) [6, 34].

The increase in inhibition, taking place after stroke, is sustained by hyperactivation of tonic—but not phasic—GABA_A_ channels[57]. We speculate that the increased tonic inhibition might be reflected in the aperiodic (power law) component of PSD rather than by its periodic features [58], more tightly linked to phasic inhibition [59]. Additionally, paired pulse stimulation protocols evidence- paradoxically- decreased GABAergic activity after stroke both in vitro [60] and in vivo [37]; this is possible since such protocols are sensitive to GABA_A_ phasic but not tonic activity[35].

A comparison between the SE and metrics derived from TMS-EEG measurements[61] which are in principle more sensitive to EEG changes related to phasic inhibition, could yield interesting insights regarding these phenomena.

Interestingly, the SE difference between the AH and UH yields values up to −0.3/−0.4 (absolute SE unit) Fig.2. These changes are in the same order of magnitude of those induced by general anesthesia with Ketamine (−0.5),[17]. Thus, stroke causes a steepening in SE that is comparable with extreme conditions such as general anesthesia.

Due to the previously mentioned limits of narrow band measures, other measures addressing the scale-free structure of EEG activity are growing in popularity, and lately methods for estimating the power-law behaviour of PSD have been implemented in popular toolboxes, such as Brainstorm[7]. Independently from other scale free measures, SE has the advantage of being simple to interpret, fast to compute and, most importantly, neurophysiologically grounded to the idea that the E/I balance shapes the overall PSD decay [24, 62]. These features make it a putative candidate for reliably assessing the state of cortical circuits following focal brain injury at the patients’ bedside.

### Potential clinical relevance of the SE

SE was positively correlated with the functional outcome of stroke patients, as assessed by the NIHSS scores, thus highlighting its potential as a predictor of the clinical features of stroke. This finding allows the discussion of some interesting aspects.

SE in all hemispheres and timepoints correlates with the effective recovery rate, ER. Notably, steeper slopes at T0 over the lesioned hemisphere were correlated with a lower degree of effective recovery, suggesting that broadband slowing in the lesioned hemisphere, will negatively affect the chances of improvements. This finding is particularly interesting and hints to a potential role of SE in predicting the functional outcome of stroke.

When assessing the link of SE with NIHSS, we notice that the correlation of SE over the affected hemisphere with the NIHSS at-T0 is not significant. This may be due to the presence of outliers that suffered stroke in less “eloquent” brain regions, thus attaining low NIHSS scores at both T0 and at T1, despite the presence of a large ischaemic lesion and a severely affected SE.

On the other hand, the SE over both hemispheres at T1 showed a good correlation with NIHSS scores at the same time point. Perhaps, in a more chronic condition, the correlation between clinical scores and EEG parameters can be better appreciated since the impact of transient factors, such as oedema[63] and post-stroke inflammation[64], that are typically present in the acute phase, is reduced with time.

Ultimately, the SE seems to be a potential marker of clinical outcome and future work should compare it with other qEEG metrics as a predictor of common clinical indexes in stroke, such as the Rankin[20] or Fugl-Meyer[22] scale. Additionally, the commercialization of dry EEG caps [65] with short setup time, will make EEG potentially viable even in acute and subacute settings, such as thrombolysis[66]. Thus, in the near future quantitative EEG measures might have a significant role in predicting clinical outcome in stroke patients and implementing personalized approaches aimed at enhancing functional recovery, like non-invasive brain stimulation techniques[67, 68] and EEG-guided robotic rehabilitation [69]. Overall, SE provided a good correlation with NIHSS.

### Cortical and subcortical stroke

We did not find significant SE or spectral differences when comparing cortical and non-cortical subgroups in our population. Recent findings by Sarasso et al [61], using TMS-EEG, showed clear differences in TMS-evoked oscillatory activity between cortical and non-cortical lesions. This could suggest that SE addresses aperiodic features that are shared by cortical and subcortical lesions, while the two conditions have differences in the frequency domain that can be better appreciated with perturbation approaches.

Previous literature reports clear spectral changes in MCA cortical strokes [19], while similar evidence is less consistent for subcortical, lacunar and posterior circulation strokes [70]. Further data on SE in various types of stroke should be collected, in order to confirm that the SE is sensitive also in cases of subcortical stroke, as it seems according to our data. In principle, while cortical lesions are known to have increased delta power, subcortical lesions do not consistently show this alteration. Some authors described reduced spontaneous EEG complexity in cases of subcortical stroke[26], which mostly relates to loss of fast activity. Reduced complexity could produce faint changes in the overall broad-band shape of PSD, that could be detected more easily by SE compared to other qEEG metrics.

### Limitations

Our study has some limitations that need to be addressed in next studies. Our sample size was relatively small and further studies on a larger population should be conducted to understand the sensitivity and specificity of SE in stroke. Patients enrolled had mild NIHSS scores. Since we needed compliance to EEG recording most severe patients had to be excluded and this might be the cause of potential selection bias. Additionally NIHSS scale has some limitations as a metric bearing relation with qEEG [71], being skewed towards the dominant hemisphere it might underestimate non-dominant hemisphere lision.

The study design did not include appropriate imaging sequences for lesion volume calculation. As such, at this stage we could not explore the relation between lesion volume and SE. Also, the low density of EEG sensors in our study does not allow for an assessment of the spatial resolution of SE with respect to the lesion location. Studying the behaviour of SE with high definition EEG could allow to study both local and long distance effects of ischemic lesions[72]. Future studies could address these important questions by employing dense array EEG setups and appropriate MRI sequences.

## Conclusions

With this study we employed the SE as a synthetic qEEG measure, reflecting the power-law decay of the PSD background, to describe the neurophysiological fingerprint of stroke. The SE appears to be a good read-out of the neurophysiological state of cortical circuits following focal ischaemic lesions. Indeed, the SE showed inter-hemispheric differences consistent across all the patients in our sample, and a partial/complete renormalization over time in most patients, which followed the amount of functional recovery, as measured by clinical scales. The use of SE in stroke is coherent with the putative neurophysiological mechanisms underpinning brain ischaemia. A simple yet effective measure as SE can be useful in the development of predictive models of outcome, or as a quick read-out for brain monitoring in ICU and could help guide future neurostimulation protocols.

---

**Supplementary Table.1 Study population.** The table shows briefly: age, affected side, stroke location and NIHSS at T0 and T1.

NIHSS=National Institute of Health Stroke Scale; R=Right; L=Left; T0=6 days after acute event(median); T1= two months after acute event (median).

**Supplementary Table.2 PSD comparisons.** Here we show the results of multiple non-parametric tests that compare power in the AH and UH at T0 and at T1, we highlight how significant differences mostly in the delta band and Delta/Alpha ratio and are significant exclusively between T0 and T1 and not between AH and UH.

AH=Affected Hemisphere; UH=Unaffected Hemisphere; T0=6 days after acute event(median); T1= two months after acute event (median).

**Supplementary Figure.1.**
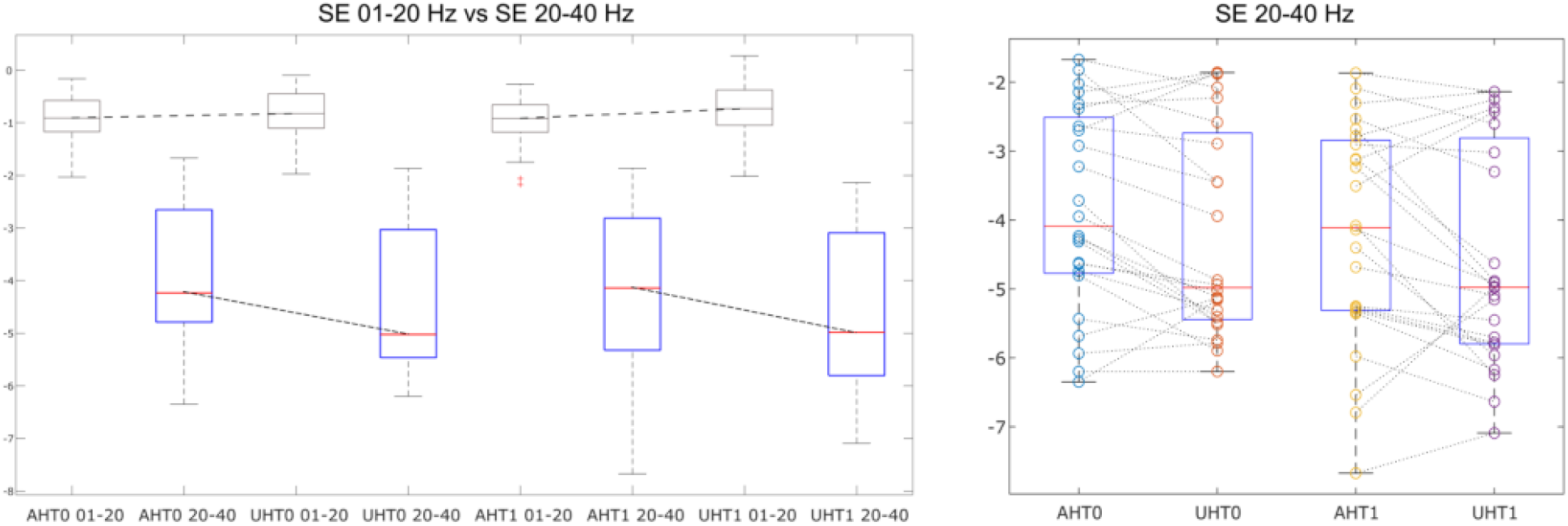
Exploring SE in the 1-20 Hz and 20-40 Hz range: In this supplementary Figure we evidence an interesting difference between the trend of SE in the 1-20 Hz range (that closely resembles the 1-40 Hz) and that of SE in the 20-40 Hz range. We speculate that, since the best results in terms of localization are provided -in our experience- by the 1-40 Hz range, the SE in the 20-40 Hz could act like a compensation factor, or rather reflect a reorganization of the neuronal time-scales following brain injury. Further studies should address this interesting topic.

## Notes

### Competing Interest Statement

The authors have declared no competing interest.

